# A method for continuously monitoring the quality of Masson pine seedlings

**DOI:** 10.1101/2020.06.01.127480

**Authors:** Li Wan, Handong Gao, Zhuo Huang, Chao Li

## Abstract

Root growth potential (RGP) is a popular physiological indicator used to evaluate seedling vigor. However, the time scale used in the RGP test is the order of days, which leads to poor performance of the RGP method. We propose an optical interference method, called statistical interferometry, to measure minute root elongation at a sub-nanometer scale, which can decrease the time used in measuring RGP. The time scale of this method is also 10^4^ times less than that of the RGP method. Because we can measure the length of root elongation continuously, we can compute the root elongation rate (RER), which is the variety of the length of root elongation per second. Continuous monitoring can help determine the quality of Masson pine seedling as soon as possible. To show the effectiveness of our proposed method, we designed an experiment, in which we applied different water stresses to our collected Masson pine seedlings and acquired two groups of pines, representing two different qualities: one stressed by water and one not. After measuring the RER of the groups in our experiments, we found that RER is interrelated with the quality of seedlings.

**Author summary:** Li Wang have worked in Nanjing Forest University. His interesting includes machine learning, computer vision and plant phenotype.

## 1 Introduction

In Southern China, Masson pine plays an important role in forest restoration, but planting it is not a trivial undertaking. At the first stage of forest restoration, intensive nursery and silviculture practices are required to ensure seedling health [1,2]. As a part of these practices, seedling quality is important in ensuring successful forest restoration. Various seedling quality assessment methods have been designed and applied, including methods to measure morphological and physiological plant attributes [2–4]. Some common seedling quality attributes are related to roots, such as root mass, shoot-to-root ratio, root electrolyte leakage and root growth potential (RGP) [5–8].

The root system is important in seedling quality assessment because roots can help a plant to uptake water. The root attributes contribute to root growth rate which affects plant growth rate and helps plants overcome planting stress [9]. Root growth has long been recognized as being important in enhancing the establishment and growth of seedlings [9–11]. With rapidly expanding root systems, seedlings can mitigate water stress [9] and quickly establish a proper water balance.

When planting a seedling, the initial root growth is critical for seedling survival and growth [12]. This attribute is assessed by the root growth capacity or potential (RGP) test, in which the seedling is kept under controlled environmental conditions and root growth is measured after a fixed time. RGP has been considered to be an indicator of a seedling’s ability to grow roots [4,13] and a measurement of a seedling’s field performance potential [2,14]. For RGP is important for a plant uptaking water and nutrients from its roots, and a high RGP can improve a plant’s chances of survival and growth [3,15].

Although RGP is an effective indicator and measurement, it requires sufficient time, usually several days. The slow growth rate of roots can increase the time to obtain the RGP because the root shapes changes dynamically and slowly, and we lack an efficient measurement method with a high spatial resolution. To solve the problem, we need a high sensitivity measurement technique that can not only obtain the RGP quickly but also monitor root growth in real time.

To increase the sensitivity and accuracy of root growth measurements, people have developed different methods based on different principles. Imaging techniques have been used to obtain the measurements through the processing of images, but their maximum attainable sensitivity is limited to the order of the wavelength due to the spatial resolution of the optical imaging systems [16-19]. The optical interference method can achieve a sub-micrometers resolution through the wavelength of laser light, and it has been used to investigate plant growth dynamics on shoots, leaves, and flower bugs with very high sensitivity and high reliability over many years [20-25]. The statistical interferometric technique [26] is also based on optical interference, but it can achieve a sub-nanometer resolution and a higher accuracy than conventional interferometric techniques.

Inspired by Rathnayake et al [27], we want to judge the quality of Masson pine seedling in less time through measuring root elongation with the statistical interferometry technique, which requires much less time than the RGP test. To show that the statistical interferometry technique can differentiate seedling quality through the root system, we applied different water stresses to two groups of Masson pine seedlings. The less water a seedling can uptake, the less energy the root can obtain through photosynthesis, which should affect root growth rate. With the statistical interferometry technique, we can monitor root elongation instantly and continuously.

Our main contribution lies in that we find the behaviors of roots are different from two different perspectives: microcosm and macrocosm. In a macrocosm, RER may always increase in a long interval. In a mincrocosm, RER changed alternately between positive and negative values in a short interval, so we cannot differentiate them simply through a threshold corresponding to RER. We propose the use of statistical features, like histograms, to represent the distribution of RER. We also propose treating an RER as a temporal signal and analyzing it through signal processing methods, like Fourier Transform. The rest of this paper is organized as follows. Section 2 introduces how we created two seedling groups and measured their RER. Section 3 gives the results we obtained in the time domain and frenquency domain. In Section 4, we explain why our proposed method is superior to the RGP method. Finally, Section 5 gives conlutions and a possible research direction.

## 2 Materials and Methods

### 2.1 Plant materials

Each Masson pine seedling was buried in a container made of a plastic envelope that is displayed in Fig 1(b). The container is a 10.5cm round full plant pot. Compared with bareroot seedlings, container seedlings have a greater root growth potential [28,29], which should benefit our ultimate goal: making the best quality seedlings. These seedlings are planted with sufficient nutrient levels and sunshine for good growth. When a seedling had grown for two years and its height was above 50cm, its roots extented along the envelope of its envelope container. Then, we could take the seedling out of the case and expose the roots for measurement, as displayed in Fig 1(a).

**Fig 1.**
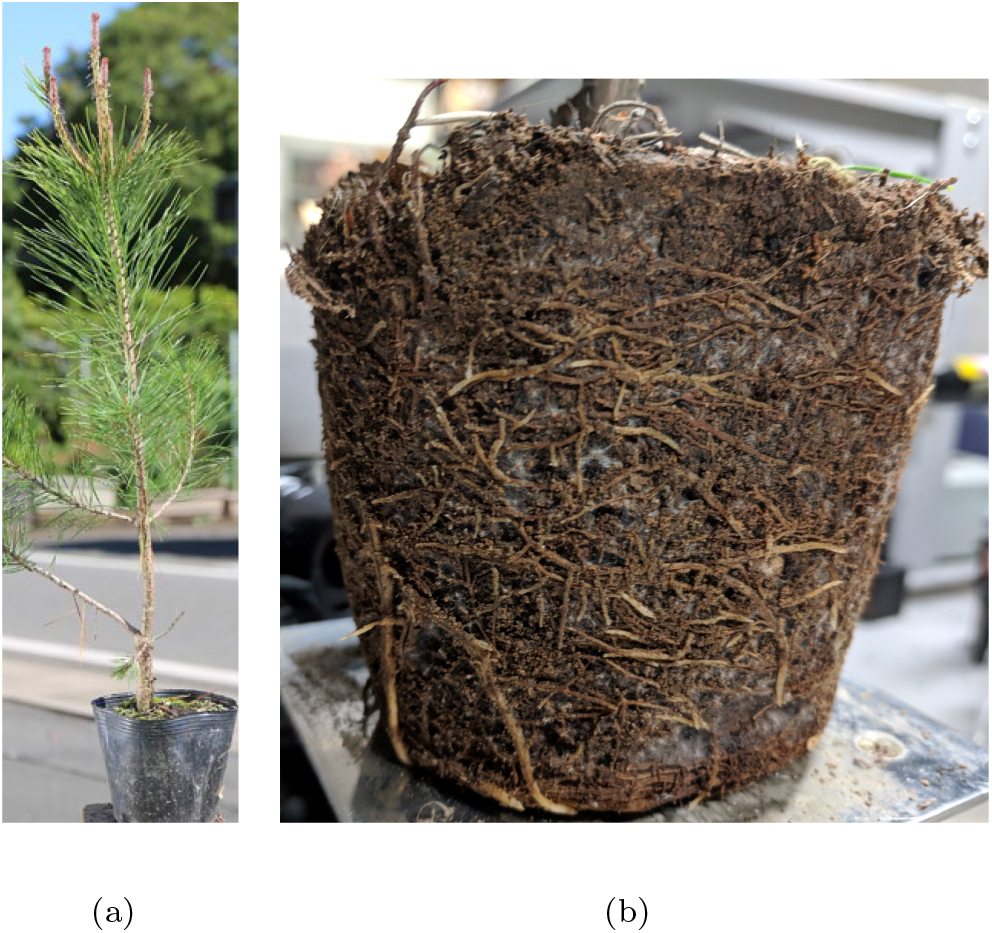
A container Masson pine seedling with its’ roots. (**a**) A container Masson pine seedling. (**b**) A Masson pine seedling’s roots..

We selected two years old seedlings due to the difficulties in exposing their roots. With less growing time, the roots could not grow enough to extend along the envelopes. To expose these roots, we would need to remove the soil from the roots by washing with water, which would definitely be a disastrous for the roots, as it will cause serious water stress to the roots and cause them to dry quickly. In contrast, if we preserved the roots in some water, the impurity and deflection of the water could disturb the projection of a laser and the camera photography, making it difficult to measure the roots’ elongation. After considering all the conditions, we decided to only take the envelope from the seedlings. The roots buried under the soil were then exposed in half of the area, which allowed them to be measured in a reasonable time and with minimum effects from the negative conditions.

### 2.2 Plant cultivation

To show that our method can differentiate betweentwo different kinds of masson seedlings: one that can absorb more water and has a high growth potential and one that can absorb less water and has a low growth potential, we applied water stress to Masson pine seedlings to simulate two different environmental conditions: dry soil and wet soil.

We moved 20 pine seedlings into a room with a mean temperature of 25 degrees. We randomly selected half of them and coved their soil with plastic sheets to preserve the water in the soil. The soil of the other seedlings was exposed to air and dried more quickly. After three days, the exposed soil appeared dry and some cracks had emerged on the surface of the soil, but the covered soil was still wet. After preparing these seedlings, we used them for the following experiments.

### 2.3 Plant root measurement

Statistical interferometry was first developed to measure the deformation of an object with a rough surface(Kadono and Toyooka, 1991). When the rough surface is illuminated by a laser light that can be randomly scattered by the rough surface, the scattered light is superposed and generates a granular pattern (speckle pattern) with high contrast. A speckle pattern generated from a root of a Masson pine seedling is shown in Fig 2.

**Fig 2.**
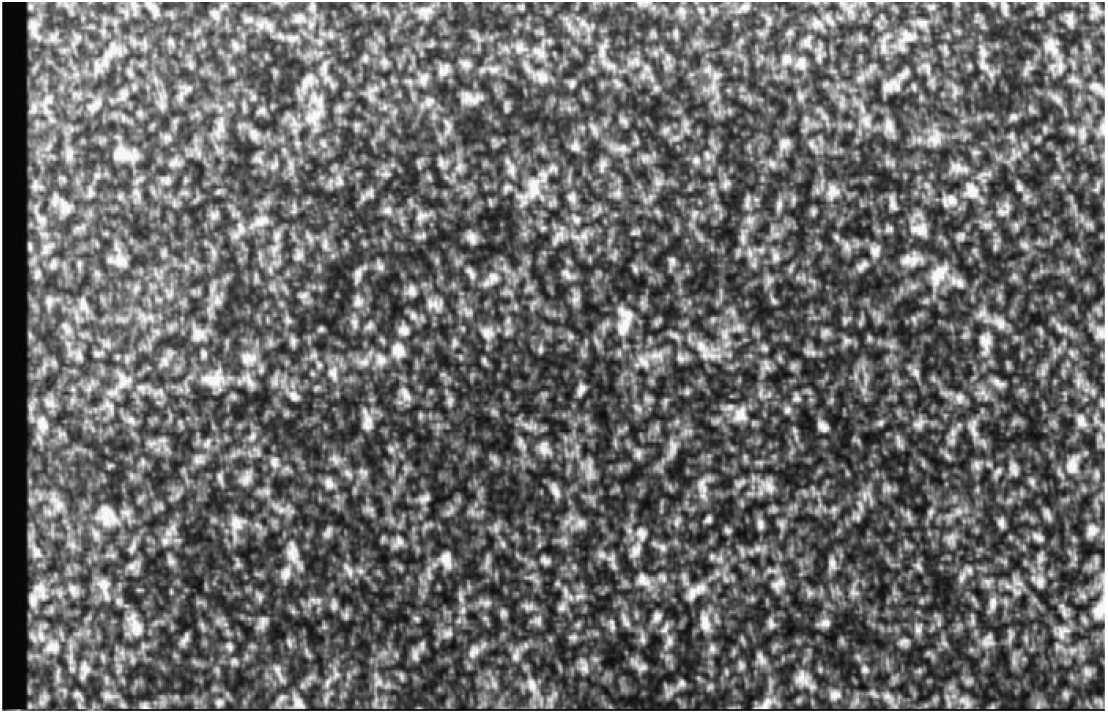
A speckle pattern generated on the illumination point on a root of a masson pine seedling.

The light field of the speckle pattern is called a fully developed speckle field. Its statistical properties are stable and independent of the scattering characteristics of the object. In this speckle field, the speckle phase is uniformly distributed from -*π* to *π* with a uniformly distributed probability density function that is used to determine the object phase through statistical interferometry [27].

The principle behind our experimental machine is displayed in Fig 3. A light emerging from an He-Ne laser(wavelength,λ = 633*nm*) is divided into two beams that illuminate the pine seedling roots. With a scattering effect, two speckle fields are generated and further superposed. The interference pattern between the two speckle fields is observed through a polarizer by a CCD camera. When the root sample elongates by Δ*x* between the two illuminating points, the optical path *L* between the two interfering speckle fields is changed by the amount Δ*L*. The relationship can be expressed by,

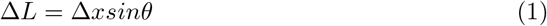

where *θ* is the angle between the illuminating beams and the observing direction. The variation in Δ*ϕ* is determined by Δ*L* according to the following equation,

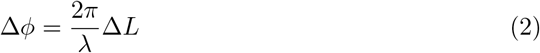

where λ is the wavelength of the laser light. After introducing equation 1 into equation 2, we obtain

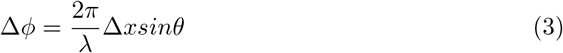

**Fig 3.**
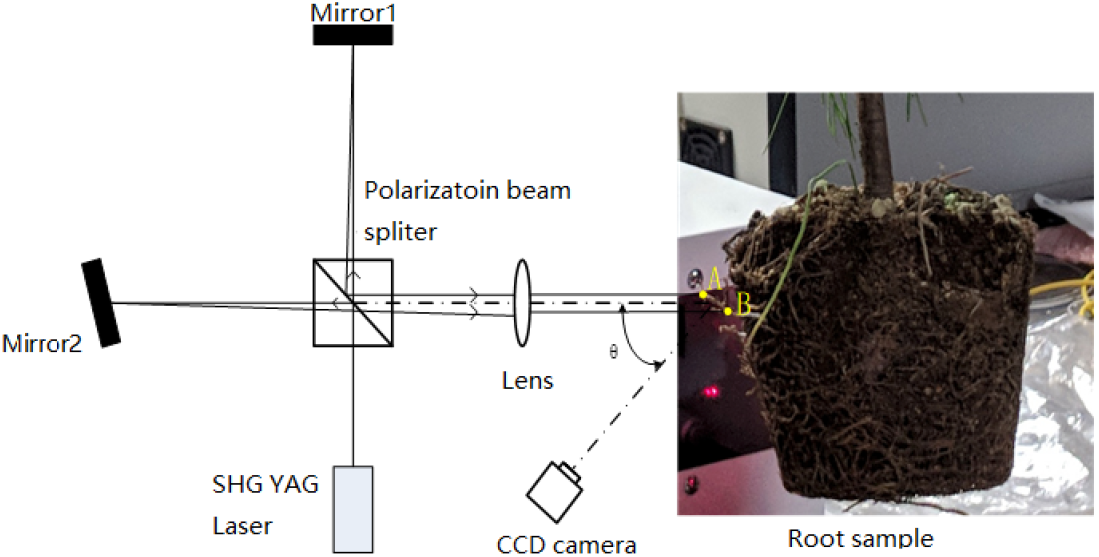
Optical system for root elongation measurements.

By analyzing the interference speckle patterns through statistical interferometry, we can obtain the speckle phase, Δ*ϕ*. Substituting it into equation 3, we can obtain the root elongation value, Δ*x*. More detailed explanations are provided in [27,30]

After preparing the machine as shown in Fig 3, we could measure the root elongation. To measure the roots, we peeled off the soft plastic case, and identified the roots without any deformation as shown in Fig 1. Most fast-growing roots can be observed on the surface of the soil, which was convenient for our experiments. We did not need to remove the soil from the roots. We only focused on the exposed roots without destroying the environments in which they lived, and the roots could survive enough for our experiments. We put each seedling on a platform that could be lifted up and down, which helped place the selected root along the optical path of the two laser light beams as shown in Fig 4. More than one root could grow along the plastic case wall, which gave us more choices. We randomly chose a lateral root with a width of 1mm for illumination. To obtain the elongation measurements of the root, we aimed for the elongation zone which was on the tip of the root [31], and we chose two points for illumination.

**Fig 4.**
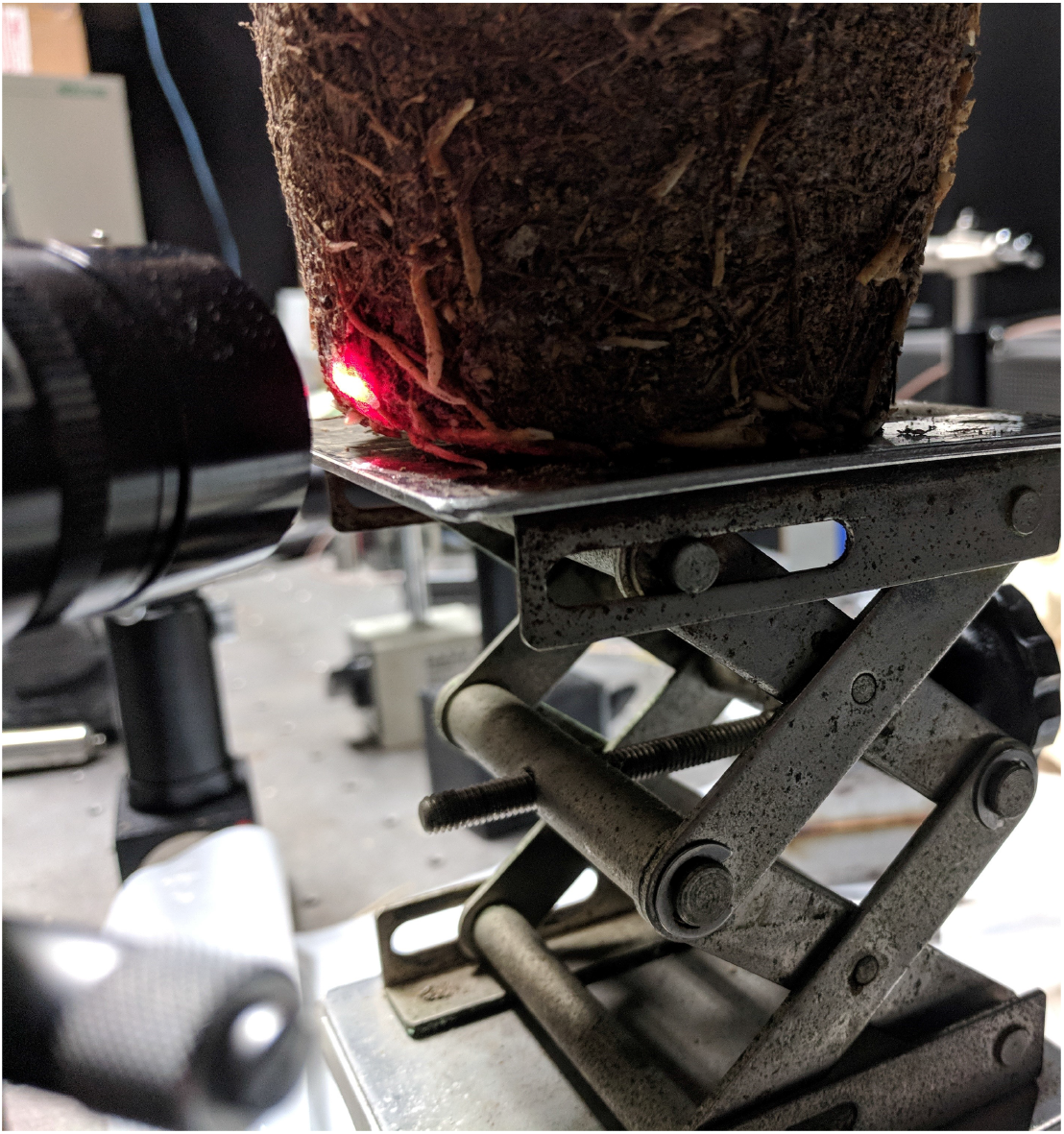
Two illuminating points are cast on the tip of a root.

Before using the experimental machine, we needed to set some parameters. The two illuminating points were divided by 3mm, so a 3-mm segment of a root sample could be monitored continuously. We measured the segment behavior with a 0.5-second interval for 150 seconds. Each root was measured at least two times. Two roots from each seedling were selected for replicate experiments, and 13 seedlings were used.

### 2.4 Statistical analysis

We used the elongation rate to distinguish the two kinds of Masson pines. The elongation rate can be plotted on the coordinate system, and the distribution of its points can help us find the difference between the two kinds of pines. Another method is through counting the frequency of the different rates that the root has exhibited. Because root growth rate is continuous, we need to divide the range of root growth rate into several intervals and compute the frequency of the rate for each root. We display the frequency in the histogram of elongation rates. Finally, we use box-and-whisker plots to show the statistical characteristics of each root. From the box-and-whisker plots, we can obtain the minimum, first quartile, median, third quartile, and maximum values for all rates.

## 3 Results

We continually experimented from 19:17 to 08:11 the following morning. The elongation rates of the roots changed dramatically between the beginning and the ending. The average rate and standard variance of the roots are displayed in Fig 5. The large variance may be caused by some conditions, such as the cycle phenomena of Masson pine seedlings. We chose the duration between 22:00 and 6:00 the following morning for our experiments to avoid the effect of some noise.

**Fig 5.**
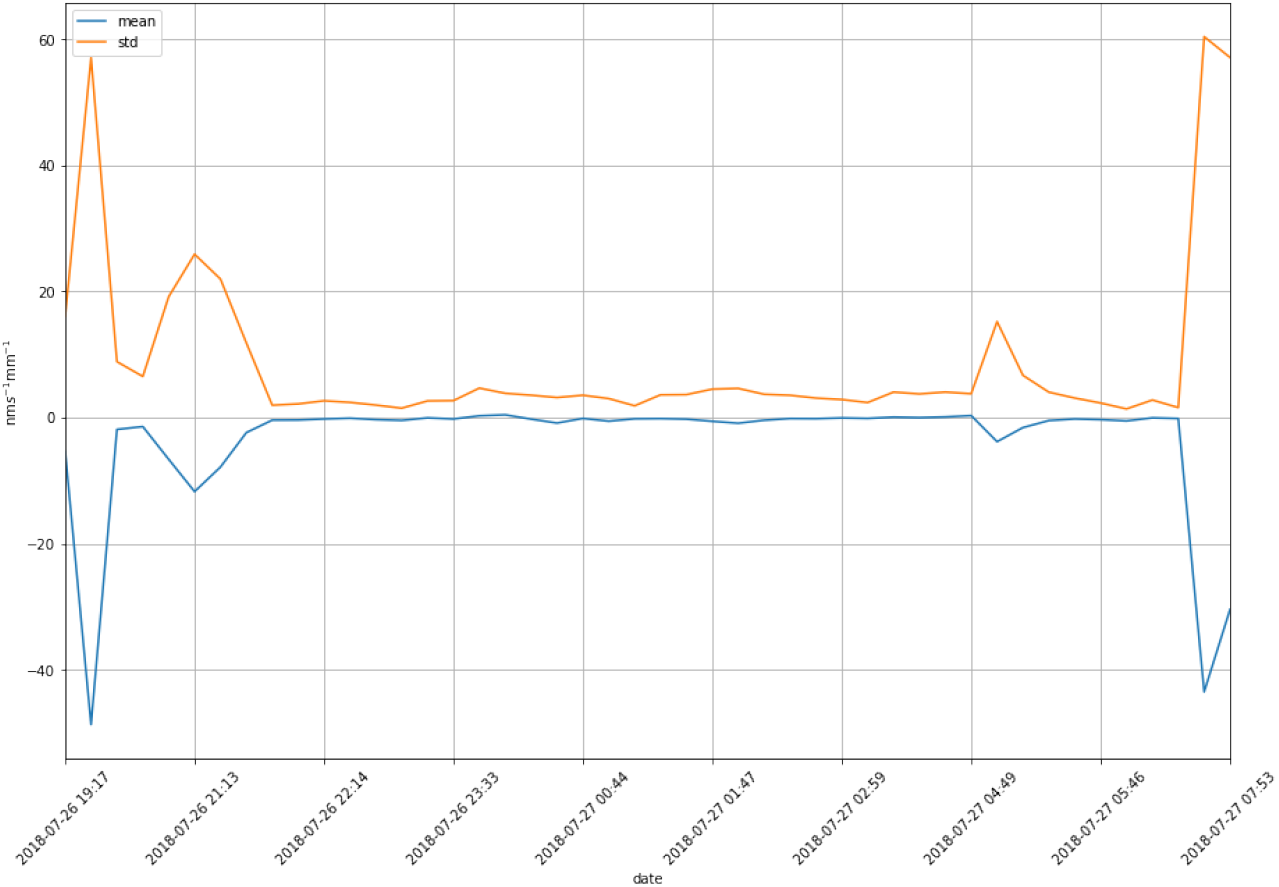
The mean and std of the elongation rates of the seedling roots.

First, we randomly select two roots to display their elongation rates. One root is from a Masson pine seedling in a water stress environment(dry soil); the other is from an environment without water stress(wet soil). The elongation rates of the roots and their statistics, including their histograms and box-and-whisker are displayed in Fig 6(a). Then, we also display the roots’ elongation rates of the entire sample and their statistics in Fig 6(b). Because we measured the 3-mm segment of a root sample in 0.5s intervals, we can compute the rate as 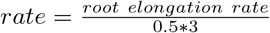, for which the unit is *nms*^−1^ *mm*^−1^.

**Fig 6.**
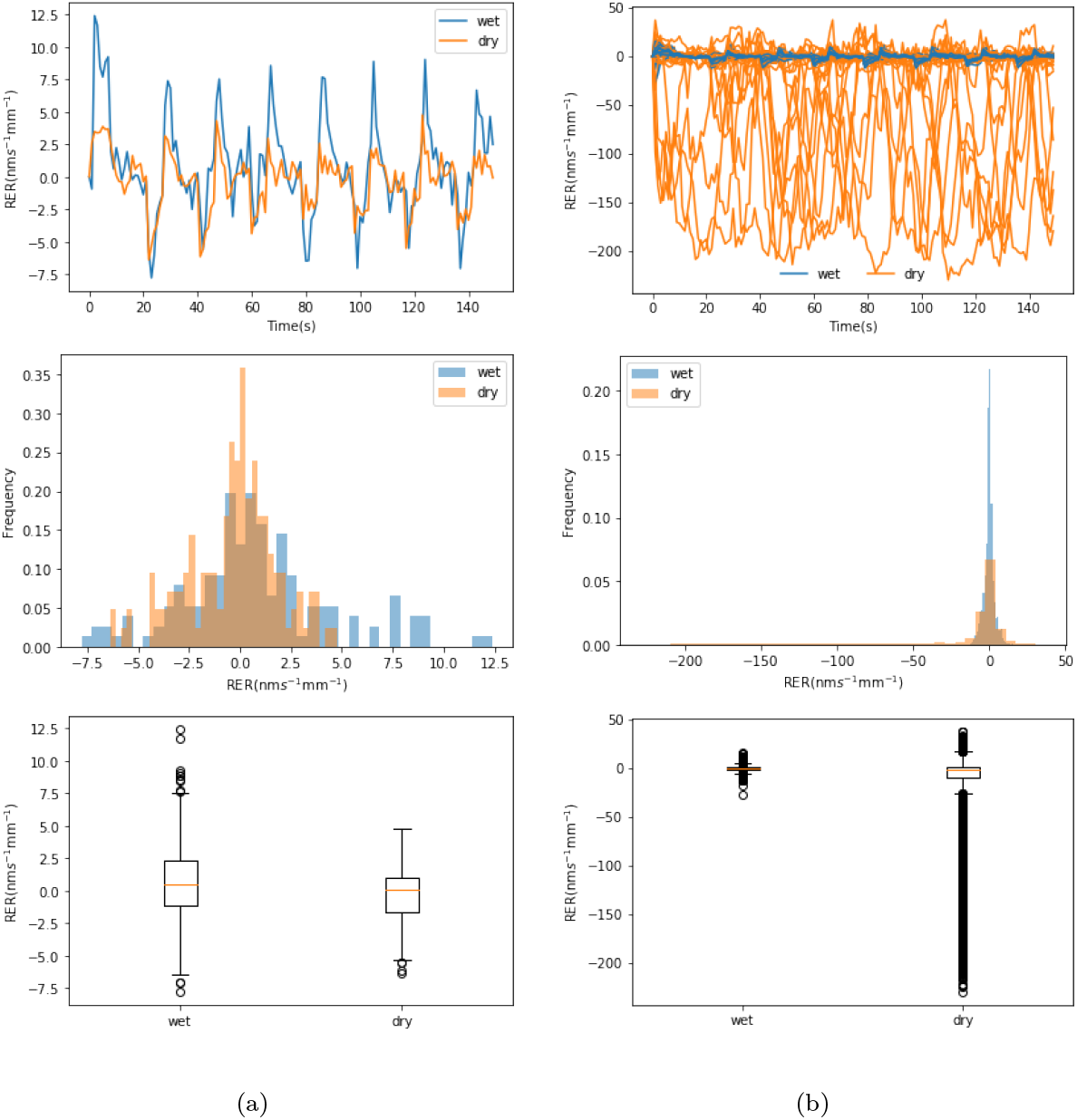
A comparison of the RER values of the roots’ in two kinds of environments. (**a**) A root elongation rate and its histogram, plus a box-and-whisker plot. (**b**) Theelongation rate of all the roots and their histogram, plus a box-and-whisker plot.

In Figure 6(a), the change in the root elongation rate(RER) in the dry soil is similar to that in the wet soil. They both behave in a periodical fashion. Most of the time, the RER in wet soil is above the RER in dry soil. In the histogram, some blue bars(wet) are on the right side of the yellow bars(dry), which means that the root length in wet soil sometimes increased while the root length in dry soil increased less and even sometimes decreased. The box-and-whisker plot also proves this. In the box-and-whisker plot, regardless of the condition the soil, the median is similar and near zero. However, most of the elongation rates in dry soil are less than zero and the amount in the lower quartile is much larger than that in the upper quartile. Most of the outliers are related to wet soil and located on the top, but the ones related to dry soil are lower, which means that the length of the roots in wet soil increased most of time; on the contrary, the root length in dry soil decreased most of the time.

We also display the RER of all root samples and their statistics in Fig 6(b). The RER of the roots in dry soil is obviously different from that of the roots in wet soil. The blue lines, which represent the RER of the roots in wet soil, are mostly lower than the yellow lines, which represent the ones in dry soil. In the histogram plot, the yellow area extends far to the left, while the blue area is mainly displayed near zero, meaning that the length of the root in wet soil may increase slightly for a short time, but the length of the root in dry soil decreases much more than that of the former. The differences between these two contexts are clear in the box-and-whisker plot in Fig 6(b). Their outliers are different: the outliers of the roots in dry soil are mainly in the negative area, while the outliers of the roots with wet soil are mainly near zero.

Interestingly, the RER seems to be cyclical, and the period is approximately 10 seconds, regardless of the environment in which the roots lived. However, the magnitude of the root’ RER in wet soil is larger than that in dry soil, which should be obvious in frequency domain if we consider RER as a temporal signal in the time domain, so we used Fast Fourier Transform (FFT) on an RER which is displayed in Fig 7(a) and obtained a result that is displayed in Fig 7(b). Although their frequency ranges are similar, less than 1.5Hz, their amplitudes are different: the amplitude with the wet group is almost double that with the dry group.

**Fig 7.**
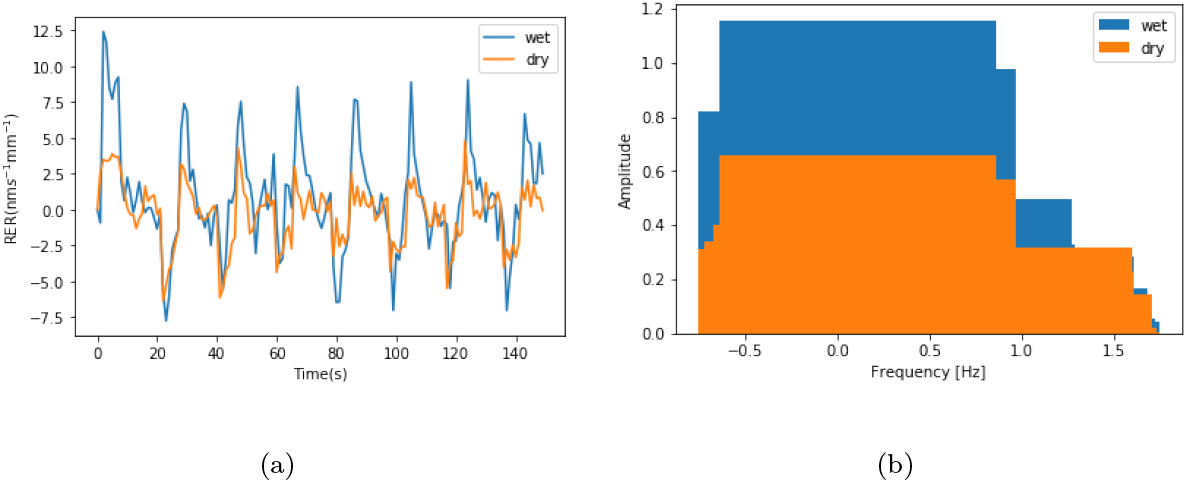
Applied Fourier Transform on RER in temporal domain. (**a**) RER in time domain (**b**) RER in frequency domain.

In addition to finding the qualitative difference between the two groups from Fig 7, we want to compute their statistical difference with statistical tests, like the Rank-wilcoxon [32], *t*-test [33] and kernel change detection [34]. To show that the wet group and the dry group have significantly different mean RER values, we apply the two sampled *t*-test to the two groups.

As the *p*-value is nearly zero in Table 1, the means of the RER values between the two groups are significantly different.

**Table 1.** The results after applying two-sampled *t*-test to the two groups of seedlings.

| group | Mean | std | $p$-value |
| --- | --- | --- | --- |
| wet | -0.10 | 3.41 | 7.38e-125 |
| dry | -23.44 | 52.63 |  |

## 4 Discussion

The RER is difficult to measure for roots that are buried under the soil. If roots are exposed to the air, they will lose water very quickly. Therefore, our experiments should satisfy two conditions.

First, the roots should not be completely deprived of soil. Because roots are likely to grow widely in soil to obtain water, they will eventually reach the wall of a container and continue growing along the wall of the container. At this time, if we peel off the wall, we can easily observe that the main body of the root is still wrapped in the soil, and only a partial surface of the root exposed to the air. Because we used the He-Ne laser(wavelength, λ), even a tiny area of a root is enough to be illuminated by the laser and the laser speckle fields can be captured by a CCD camera clearly as shown in Fig 2. The two illuminating points are also 3 mm apart. Thus, we can easily select an optional root to measure.

Second, the measurement time should not be too long. Even if we expose a partial area of a root, we unavoidably change the environment in which the root lives. Complex environments may introduce unpredictable effects. To avoid most environmental effects, we should decrease the measurement time. Fortunately, our method can implement macro-scale measurements. Even if the measurement time is decreased to several minutes, we can still obtain the status of root growth, including the cycle time of the root elongation rate, which is approximately 10s.

After satisfying the above two conditions, we expect to see that roots in wet soil will grow faster than roots stressed by little water. Water is necessary for photosynthesis, which can provide some energy for root growth. Roots with more energy can grow more quickly, and subsequently uptake more water from more soil and, in turn, strengthen the photosynthesis process.

Based on photosynthesis theory and root function, we think that RER reflects the vigor of a seedling. RGP can be treated as a method for measuring RER. The weakness of this method is that it requires ten or more days for some roots to grow to a visible size and be measured by traditional tools, such as a ruler. However, RER can be measured with our method after several minutes.

There is a substantial difference between our method and the RGP method. In the RGP method, roots will always grow larger after a long time if the living conditions are good, but RER does not always increase in a short time. In Figure 6(a), the RER fluctuates around zero; it is negative at some times and positive at other times. This means that the root will occasionally shrink regardless of the conditions the seedling lives in. In [27], the measurement was performed in 0.5s time intervals for 3.5s and the mean value is computed. The same measurement continued for a period of 9.5 min in 30s intervals. The experiments show that root length will always increase and that the RER will always be positive. In fact, we think that only using the mean value will cause some information, like the cycle time and the magnitude of a signal, to be lost. Based on our results, we can obtain all the information without losing these piceces of data.

The RER is varied during a short time period and does not remain constant, so we cannot distinguish seedlings only through their RER values. Further, the RERs of seedlings living in two different conditions overlap over time. We think the statistical calculation will help us distinguish between these different seedlings. In Figure 6(a), we plot the RER variations along the time axes. The RER values related to wet soil changed more than those related to dry soil. Surprisingly, the roots in wet soil shrank more than those in dry soil, which seems to violate our intuition. However, the roots in wet soil also had larger RER values than that in dry soil. We think the phenomenon appropriately proves that roots without water stress have more vigor than those with water stress. In the box-and-whisker plot, the difference is also obvious based on the time domain plotted in the top row of Figure 6(a).

The histograms in Figure 6(a) exhibit fewer differences because they seems overlap most of the time. Regardless, we think that the histograms should be an appropriate basis for distinguishing the different vigor of seedlings.

When all RER values are plotted in Figure 6(b), it is a little confusing at first, especially in the top row of Figure 6(b). It seems that many RER values related to dry soil are larger than the ones related to wet soil. However, from the bottom row of Figure 6(b), we can see that most RER values related to dry soil are negative and much lower than the ones related to wet soil. From the middle row of Figure 6(b), the RER distributions related to the two conditions are different, that can be approved further by the *p*-value(7.38e-125) in Table 1.

Because the RER changed cyclically and dramatically along the time, the RER distributions related to different conditions may be overlapped and cannot be distinguished easily. If we treat RER variations as a signal in the time domain and apply FFT to them, we can capture the feature of them. The feature should help us classify the RER values corresponding to two conditions, which can be observed in Figure 7.

## 5 Conclusion

In this study, we show that our method can measure the RER of Masson pine seedlings on the scale of minutes, which is far less time than the RGP method, which requires days. Our method is non-destructive and does not involve contact and it can be used to continuously monitor the progress of root growth. Because we need only a tiny root and a short time to measure RER, seedlings experience only slight environmental effects, ensuring that we can obtain more accurate information about the roots. On the scale of seconds, we observed that the RER of roots will not always increase. Many things are different in a microcosm and a macrocosm. We can take advantage of this to provide clues about the quality of Masson pine seedlings. In the future, we can take advantage of more signal processing techniques, such as frequency spectrum, to extract the feature of RER, which should help us observe the behavior of seedlings living in different environments through a different domain.

## 6 Acknowledgement

The authors acknowledge the grant support provided by the National Natural Science Foundation of China (NSFC: 31570714) and Jiangsu Overseas Research & Training Program for University Prominent Young & Middle-aged Teachers and Presidents Program.

The authors also thank to the generosity of Kadono, Hirofumi who lend us a new machine to measure RER and their laboratory to do experiments.

## References

1. Gladstone WT, Ledig FT. Reducing pressure on natural forests through high-yield forestry. Forest Ecology and Management. 1990;35:69–78.

2. Grossnickle SC. Ecophysiology of northern spruce species: the performance of planted seedlings. NRC Research Press; 2000.

3. Burdett A. Quality control in the production of forest planting stock. The Forestry Chronicle. 1983;59(3):132–138.

4. Ritchie G. Assessing seedling quality. In: Forestry nursery manual: production of bareroot seedlings. Springer; 1984. p. 243–259.

5. Mohammed GH. The status and future of stock quality testing. New Forests. 1997;13(1-3):491–514.

6. Haase DL. Understanding forest seedling quality: measurements and interpretation. Tree Planters’ Notes. 2008;52(2):24–30.

7. Landis TD, Dumroese RK, Haase DL. The container tree nursery manual: Volume 7, Seedling processing, storage, and outplanting. Agric Handbook No 674 Washington, DC: US Department of Agriculture, Forest Service 199 p. 2010;674(7).

8. Thompson BE. Seedling morphological evaluation: what you can tell by looking. 1985;.

9. Stone EC. Poor survival and the physiological condition of planting stock. Forest Science. 1955;1(2):90–94.

10. Toumey JW, Korstian CF, et al. Seeding and planting in the practice of forestry. Seeding and planting in the practice of forestry. 1942;(3rd ed.).

11. Rudolf PO. Why forest plantations fail. Journal of Forestry. 1939;37(5):377–383.

12. Simpson DG, Ritchie GA. Does RGP predict field performance? A debate. New Forests. 1997;13(1-3):253–277.

13. Ritchie GA, Tanaka Y. Root growth potential and the target seedling. In: Target seedling symposium, GTR: RM-200, USDA Forest Service; 1990. p. 37–51.

14. Burdett A. Understanding root growth capacity: theoretical considerations in assessing planting stock quality by means of root growth tests. Canadian Journal of Forest Research. 1987;17(8):768–775.

15. Johnson JD, Cline ML. Seedling quality of southern pines. In: Forest regeneration manual. Springer; 1991. p. 143–159.

16. Schmundt D, Stitt M, Jahne B, Schurr U. Quantitative analysis of the local rates of growth of dicot leaves at a high temporal and spatial resolution, using image sequence analysis. Plant Journal. 1998;16(4):505–514.

17. Beemster GTS, Baskin TI. Analysis of Cell Division and Elongation Underlying the Developmental Acceleration of Root Growth in Arabidopsis thaliana. Plant Physiology. 1998;116(4):1515–1526.

18. Der Weele CMV, Jiang HS, Palaniappan KK, Ivanov VB, Palaniappan K, Baskin TI. A New Algorithm for Computational Image Analysis of Deformable Motion at High Spatial and Temporal Resolution Applied to Root Growth. Roughly Uniform Elongation in the Meristem and Also, after an Abrupt Acceleration, in the Elongation Zone. Plant Physiology. 2003;132(3):1138–1148.

19. Walter A, Spies H, Terjung S, Kusters R, Kirchgesner N, Schurr U. Spatio-temporal dynamics of expansion growth in roots: automatic quantification of diurnal course and temperature response by digital image sequence processing. Journal of Experimental Botany. 2002;53(369):689–698.

20. Fox MD, Puffer LG. Holographic Interferometric Measurement of Motions in Mature Plants. Plant Physiology. 1977;60(1):30–33.

21. Briers JD. The Measurement of Plant Elongation Rates by Means of Holographic Interferometry: Possibilities and Limitations. Journal of Experimental Botany. 1977;28(2):493–506.

22. Briers JD. Speckle Fluctuations as a Screening Test in the Holographic Measurement of Plant Motion and Growth. Journal of Experimental Botany. 1978;29(2):395–399.

23. Jiang Z, Staude W. An Interferometric Method for Plant Growth Measurements. Journal of Experimental Botany. 1989;40(10):1169–1173.

24. Dyrseth AA. Measurement of plant movement in young and mature plants using electronic speckle pattern interferometry. Applied Optics. 1996;35(19):3695–3701.

25. Madjarova VD, Toyooka S, Nagasawa H, Kadono H. Blooming Processes in Flowers Studied By Dynamic Electronic Speckle Pattern Interferometry (DESPI). Optical Review. 2003;10(5):370–374.

26. Kadono H, Takahashi G, Toyooka S. Monitoring of biological activity of plant using difference-image of biospeckle. 19th Congress of the International Commission for Optics: Optics for the Quality of Life. 2004;4829:988. doi:10.1117/12.527504.

27. Rathnayake AP, Kadono H, Toyooka S, Miwa M. A novel optical interference technique to measure minute root elongations of Japanese red pine (Pinus densiflora Seibold & Zucc.) seedlings infected with ectomycorrhizal fungi. Environmental and Experimental Botany. 2008;64(3):314–321. doi:10.1016/j.envexpbot.2008.02.007.

28. South DB, Harris SW, Barnett JP, Hainds MJ, Gjerstad DH. Effect of container type and seedling size on survival and early height growth of Pinus palustris seedlings in Alabama, U.S.A. Forest Ecology and Management. 2005;204(2):385–398.

29. Grossnickle SC, Elkassaby YA. Bareroot versus container stocktypes: a performance comparison. New Forests. 2016;47(1):1–51.

30. Kadono H, Toyooka S. Statistical interferometry based on the statistics of speckle phase. Optics Letters. 1991;16(12):883–885.

31. Lazar T. Taiz, L. and Zeiger, E. Plant physiology. 3rd edn. Annals of Botany. 2003;91(6):750–751.

32. Palumbo F, Aceto G, Botta A, Ciuonzo D, Persico V, Pescapè A. Characterizing Cloud-to-user Latency as perceived by AWS and Azure Users spread over the Globe; 2019.

33. Ciuonzo D, Carotenuto V, De Maio A. On Multiple Covariance Equality Testing with Application to SAR Change Detection. IEEE Transactions on Signal Processing. 2017;65(19):5078–5091.

34. Desobry F, Davy M, Doncarli C. An online kernel change detection algorithm. IEEE Transactions on Signal Processing. 2005;53(8):2961–2974.

